# Neurodesign: Optimal Experimental Designs for Task fMRI

**DOI:** 10.1101/119594

**Authors:** Joke Durnez, Ross Blair, Russell A. Poldrack

## Abstract

A recent stream of alarmist publications has questioned the validity of published neuroimaging findings. As a consequence, fMRI teams worldwide have been encouraged to increase their sample sizes to reach higher power and thus increase the positive predictive value of their findings. However, an often-overlooked factor influencing power is the experimental design: by choosing the appropriate experimental design, the statistical power of a study can be increased within subjects. By optimizing the order and timing of the stimuli, power can be gained at no extra cost. To facilitate design optimization, we created a *python* package and web-based tool called Neurodesign to maximize the detection power or estimation efficiency within subjects, while controlling for psychological factors such as the predictability of the design. We implemented both a simulation-based optimisation, as well as an optimisation using the genetic algorithm, introduced by Wager and Nichols (2003) and further improved by Kao et al. (2009), to optimize the experimental design. The toolbox *Neurodesign* allows more complex experimental setups than existing toolboxes, while the GUI provides a more user-friendly experience. The toolbox is accessible online at www.neuropowertools.org.

## 2 Introduction

A recent stream of alarmist publications has questioned the validity of published neuroimaging findings (Eklund et al., 2016; Ioannidis, 2005; Open Science Collaboration, 2015). At the core of the reproducibility crisis is the lack of power typically observed in neuroimaging (Button et al., 2013), and more specifically, fMRI studies (Durnez et al., 2014). The signal measured in fMRI is known to be very noisy, while the hypothesised effects are small, such that a push for larger sample sizes promises a more powerful future for neuroimaging. Different power analysis strategies offer a way to optimise the sample size for a specific power level (Durnez et al., 2014; Mumford and Nichols, 2008; Hayasaka et al., 2007; Durnez et al., 2016). However, fMRI data are typically acquired and aggregated on two levels: within and between subjects. As such, increasing the power of an fMRI experiment can be achieved by increasing the number of subjects, but also via the within-subjects experimental design. This is especially true for smaller and more subtle effects, where the power curve is characterised by a slower increase, and thus the resulting power is more affected by the number of subjects and the number of time points. In addition to the duration of the experiment for each subject, the order and timing of different conditions within the experiment also influence the power of the resulting analyses.

The goal in task fMRI experiments is often one of two: detection or estimation. Detection refers to detecting the difference in brain activation between conditions or groups, while estimation relates to estimating the exact shape of the evoked fMRI response (called the haemodynamic response function, HRF). Ideally, the design of an fMRI experiment changes according to the specific research question asked. An optimal design with respect to these two distinct research questions are said to maximize the detection power or the estimation efficiency respectively. It is often argued that those two goals are opposite and an increase in detection power inevitably leads to a decrease of estimation efficiency. For example, when two trials of the same condition follow each other closely, the signal tends to accumulate linearly (Dale, 1999), which makes it easier to detect. Therefore, the experiments often consist of blocks of the same condition. This type of design is called a blocked design. On the contrary, the accumulation (and saturation) of the measured signal conceals the shape of the HRF. To estimate the HRF, scientists often opt for an event-related design, where both the timing and order of conditions are randomised. However, Kao et al. (2009) show that the necessary trade-off between detection and estimation can be improved using certain optimisation algorithms.

Another important aspect in an fMRI design is the psychological experience of the subject in the scanner. With a blocked design, the design becomes very predictable for subjects which can potentially bias the psychological function under investigation. To minimise predictability, Buracas and Boynton (2002) propose the use of m-sequences. However, the length of m-sequence is restricted to *n* = (*Q*_1_)^*l*^ − 1 with *Q* + 1 a prime, *Q* the total number of stimulus types, and *l* a positive nonzero integer. Recently Lin et al. (2007) proposed the use of a circulant (almost-)orthogonal array to expand the range of possible fMRI designs while ensuring complete independence between a trial and its successor. Very often, the best design is a combination of a maximal signal with low predictability. Therefore, Wager and Nichols (2003) suggest the use of a genetic algorithm to find an optimisation between estimation efficiency, detection power and predictability. This algorithm optimises a weighted average of different criteria, with the weights depending on the hypothesis and the expected outcome of the experiment. Subsequent work has further fine-tuned the algorithm and compared it with other approaches (Kao et al., 2009). However, in some cases the design requires more control than is offered in the genetic algorithm, in which case a simulation-based optimisation is the only considerable option. In this paper, we present *Neurodesign* for fMRI design optimisation with different optimisation algorithms, which is both available as a python module as well as a GUI web tool, available at www.neuropowertools.org. The paper is structured as follows: we start with a general description of the methodology in section 2. We show how designs can be compared and optimised using our python module in section 3. An overview of the GUI is given in section 4. We compare our toolbox with other existing software in section 5 and compare the designs from different optimisers in section 6. Finally, we conclude and discuss the results in section 7 and 8.

## 3 Design optimisation using the genetic algorithm

### 3.1 Statistical measures of design optimality

The signal measured using fMRI is the blood oxygen level dependent (BOLD) signal, which is a assumed to be related to the neural signal via convolution with a hemodynamic response function (HRF). We consider the general linear model as the underlying model for the statistical objective as in Equation 1.

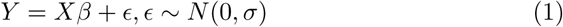

We denote *Y* as the measured signal. *X* represents the design matrix, *β* is the response amplitude for each column/condition in *X* and *ϵ* the error.

There are two types of design matrices: the convolved model and the finite impulse response (FIR) model where both are transformations of the matrix *X*_base_, a *m × t* matrix, where *m* is the number of stimuli and *t* the number of measured timepoints. The values in *X*_base_ are 1 or 0: 1 when stimulus *M* is shown on time point *T*. The two transformations of *X* are The two possible transformations are shown in Figure 1. In the first model, the regressor *X*_base_ is convolved with the hemodynamic resonse function (Figure 1, panel 2) to represent the expected signal for a brain region related to the stimulus. The second model aims to estimate the exact shape of the HRF, by including regressors identical to the stimulus presentation, but each regressor with a certain temporal lag (Figure 1, panel 3).

**Figure 1:**
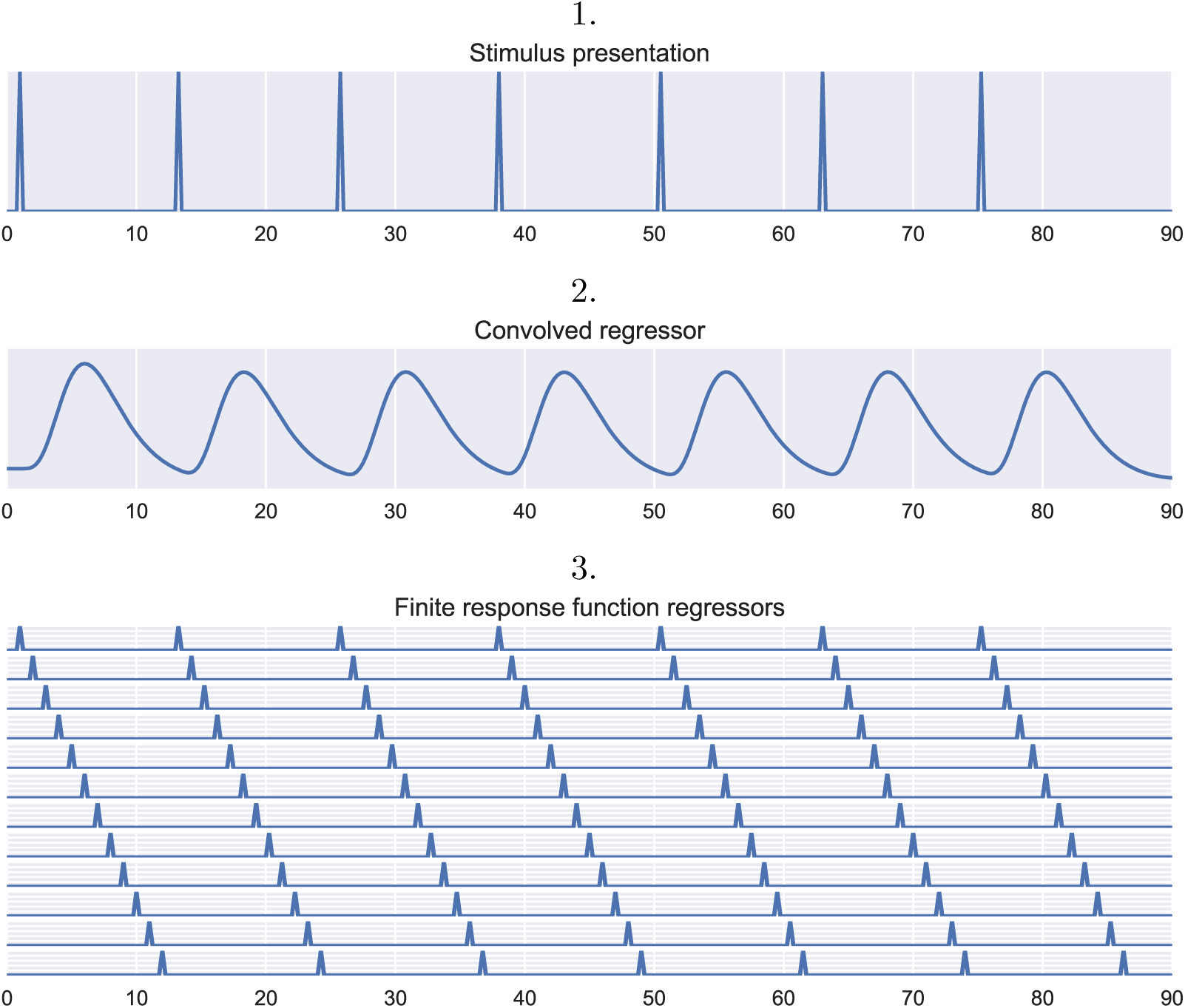
An experimental fMRI design with one stimulus type and two common models used in the GLM when modeling the resulting BOLD signal. The first panel shows the timeseries of the stimulus onsets. The second panel shows the stimulus onsets convolved with the double-gamma HRF, which can be interpreted as the expected BOLD signal if the measurement is related to the task. The parameter *β* in equation 1 with this model represents the amplitude of the signal related to the task.The third panel shows the FIR model, with each regressor a shifted version of the stimulus onsets. The *β*-parameters represent the amplitude of the HRF at specific time points following stimulus onset. Units on the x-axis are seconds. Units on the y-axis are removed, as these are meaningless and often rescaled to have unit height.

Often, researchers are interested in specific hypotheses concerning particular combinations of parameters. The parameter of interest *β* can be estimated using the least squares estimator in Equation 2 and its variance in Equation 3.

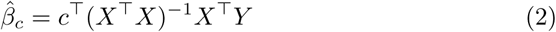

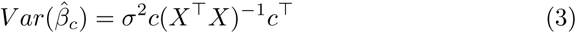

with *c* the contrast vector of interest. To account for the specific character of fMRI data, we alter the model slightly. Because fMRI timeseries data exhibit substantial temporal autocorrelation, the errors in equation 1 are not independent, *ϵ V*, where off-diagonal values represent the correlation between measurements at different time points. Furthermore a regressor, *S*, representing low-frequency noise components is added to the model (see Kao et al. (2009) for detailed derivations). The resulting variance of the estimator in defined in Equation 4.

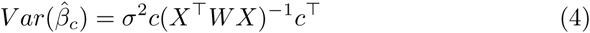

with *W* = *V* − *V S*^T^(*SV S*^T^)^−1^*SV*. An optimal experimental design with respect to the estimator minimises the variance of the estimator. We will therefore quantify the optimality of the design as the inverse of *c*(*X*^T^*WX*)^−1^*c*^T^. Most often, an fMRI experiment has multiple contrasts of interest, thus *c*(*X*^T^*WX*)^−1^*c*^T^ becomes a square matrix. With *r*_*c*_ the number of contrasts, there are two common ways to quantify the optimality of the design as defined in Equation 5.

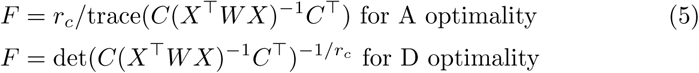

We denote *F*_*e*_ as the estimation efficiency if *X* is a FIR, and *F*_*d*_ as the detection power if *X* is a convolved design matrix.

### 3.2 Psychological measures of design optimality

Apart from the statistical concept of design efficiency, it is important to account for psychological factors that might render the experimental design invalid. The most important factor is predictability. For example in experiments addressing cognitive control, such as a stop-signal task, it is of the utmost importance that the trial type on any given trial cannot be easily predicted from the trial type on the previous trial, to avoid psychological confounding of the experiment. We quantify the optimality of the design in terms of confounding in Equation 6.

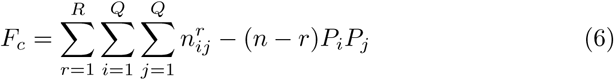

where 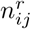 is the number of trials of type *i* at timepoint *t* preceding a trial of type *j* at timepoint *t*+*r*. The variable *P*_*i*_ is the proportion that trial should occur in the experiment. If *F*_*c*_ = 0, there are no unforeseen contingencies between trial types. The final optimality criterion controls the desired trial type frequencies: 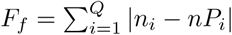, with *n*_*i*_ the number of trials of type *i*.

### 3.3 Multi-objective criterion

To ensure comparability across different optimality criteria, we first rescale the different optimality criterion to a scale of 0 to 1 as in Kao et al. (2009). To find the maximum *F*_*d*_ and *F*_*e*_ possible, we first run an optimisation with weights 1 for respectively *F*_*d*_ and *F*_*e*_ and weights 0 for the other optimality criteria. In the multi-objective criterion, the *F*_*d*_ and *F*_*e*_ scores are divided by their respective maximum to ensure scores between 0 and 1. For *F*_*f*_ and *F*_*c*_, the score for the worst possible design (a design with only the least probable stimulus) is taken as the maximum score. Second, whereas larger *F*_*e*_ and *F*_*d*_ represent better design, the opposite is true for *F*_*c*_ and *F*_*f*_. Therefore the scores for *F*_*c*_ and *F*_*f*_ are subtracted from 1. As such, the resulting optimality criteria is obtained in Equation 7.

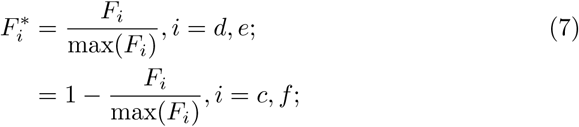

As no design can ensure optimality in all four optimality criteria, the goal of any design optimisation depends on the researcher’s goal of the experiment. Given prespecified weights *w*_*i*_ with *i* = *c, d, e, f*, Σ_*i*_ *w*_*i*_ = 1, *w*_*i*_ ≥ 0, we define the weighted optimality criterion in Equation 8.

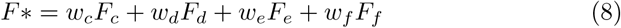

### 3.4 Optimisation algorithms

#### 3.4.1 Genetic algorithm

A genetic algorithm is a method for solving optimisation problems inspired by natural selection in biological evolution. Contrary to classical optimisation algorithms, a genetic algorithm generates a population of points at each iteration. A graphic representation of the genetic algorithm with an fMRI example is shown in Figure 2. The steps of the genetic algorithm are.

**Figure 2:**
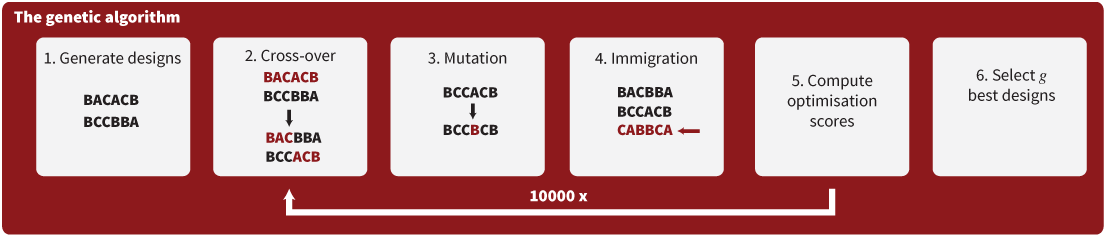
Graphical representation of the genetic algorithm. The examples in each step are pieces of experimental designs with 3 different trial types (A, B, C). In the example, the inter-trial interval is ignored.

1. Create G initial random designs.
2. **Crossover.** Pair the best G/2 designs with each other.
3. **Mutation.** Randomly switch q% of all trials by random trial types.
4. **Immigration.** Add new random designs to the population.
5. **Natural selection.** Compute optimality scores and select G best designs
6. Repeat step 2-5 until a stopping rule is met.

A crucial part of the algorithm is drawing random designs from the population of designs (in step 1 and step 4). This could for example be achieved by using *m*-sequences to decide the order of the stimuli, the stimuli evenly spaced in time. We show in section 3.4 how to sample random designs using *neurodesign*.

#### 3.4.2 Simulation-based optimisation

While the genetic algorithm has been shown to be a powerful algorithm for experimental design optimisation (Kao et al., 2009), there are certain instances where more strict control of the design is desired. For example, in cognitive control, the occurence of one condition versus another is sometimes varied between subjects. In those instances, it is crucial that there is strict control of the proportion of trial types that are shown. The genetic algorithm punishes deviance from the proportions measured by *F*_*f*_, but strict control cannot be ensured. Therefore, we included a random design generator. The implementation is identical to the *Genetic algorithm*, with the *Crossover* - and *Mutation*-steps are removed and only the *Immigration*-step is performed.

## 4 Neurodesign, python module

### 4.1 Installation

NeuroDesign is available on *PyPi* and can be installed as:

~~~
pip install neurodesign
~~~

Next, we will give an introduction to the python module. For all functionality, please refer to the manual.

### 4.2 Specifying the characteristics of the experiment

In a first step, the experiment should be described in the class called *experiment*. This contains general information, such as the number of stimuli and the duration of the experiment, but also more specific information, such as the model with which the inter trial intervals (ITI) are sampled. This function will generate the assumed covariance matrix, the drift function and the whitening matrix. All parameters are described in Table 1, while a graphical representation of components of an experiment are described in Figure 3. We define a simple experimental setup with 20 trials and 3 conditions, which we will use to exemplify the next functions:

~~~
import neurodesign
EXP = neurodesign.experiment(
    TR=1.2,
    n_trials=20,
    P = [0.3,0.3,0.4],
    C = [[1,−1,0],[0,1,−1]],
    n_stimuli = 3,
    rho = 0.3, stim_duration=1,
    ITImodel = ‘uniform’,
    ITImin = 2,
    ITImax=4
    )
~~~

**Table 1:**
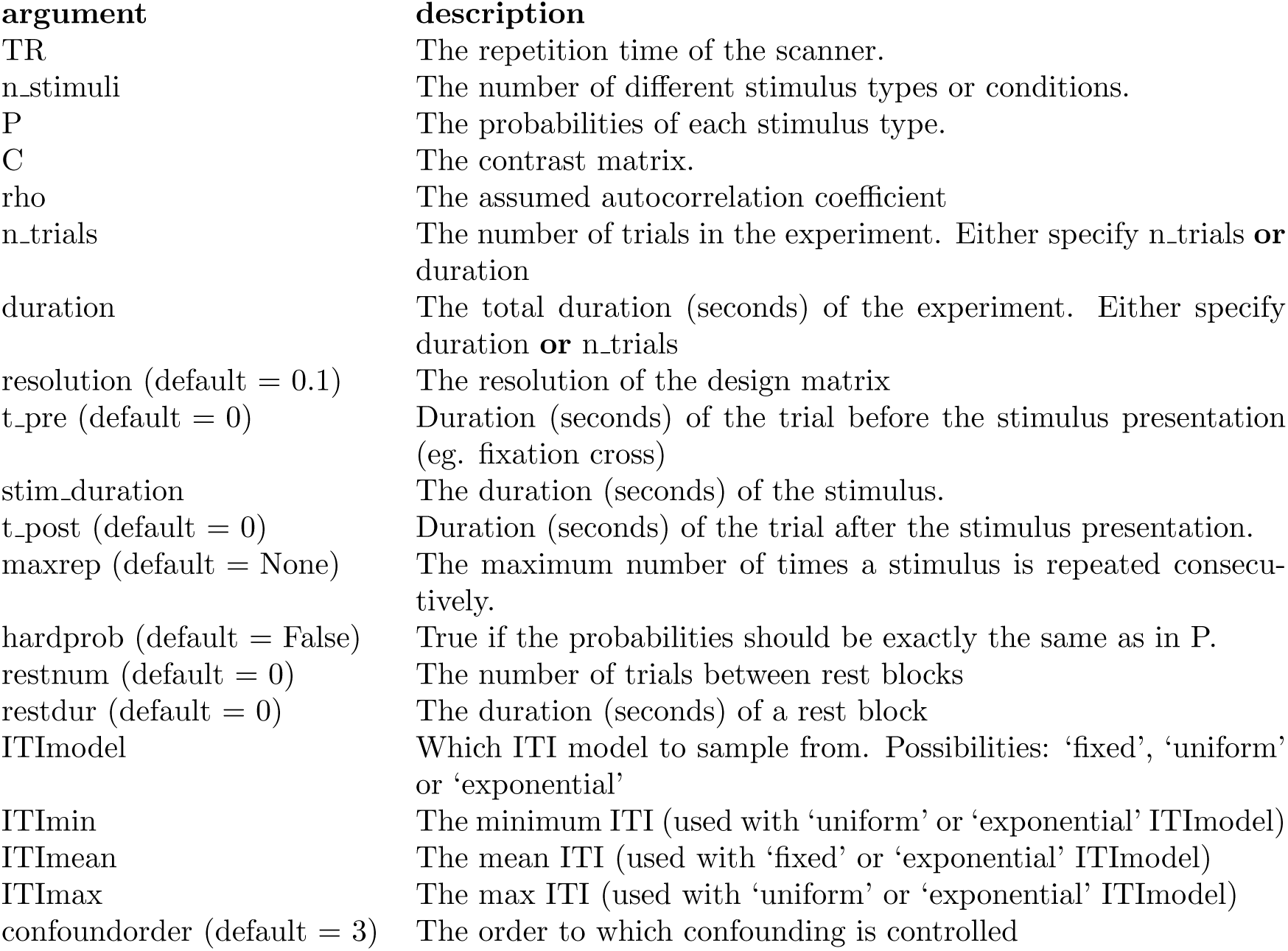
The arguments for object of class *neurodesign.experiment*

**Figure 3:**
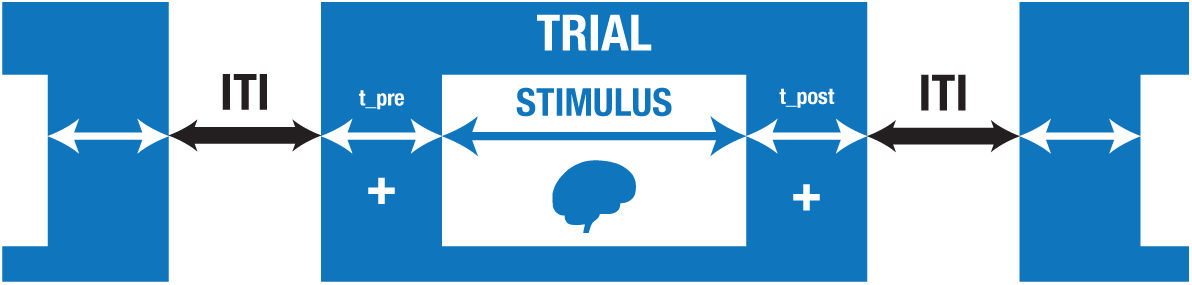
The basic layout of an experimental trial.

### 4.3 Generating a design matrix

Within the defined experimental setup, we can now define a design matrix, develop the design matrix and compute the optimality scores using the class *design*. We use equal weights for the different optimality criteria for the weighted average optimality attribute. The only input required is the stimulus order, the ITI’s and an object of class *neurodesign.experiment*:

~~~
import neurodesign
DES1 = neurodesign.design(
order = [0,1,2,0,1,2,0,1,2,0,1,2,0,1,2,0,1,2,0,1],
ITI = [2]*20,
experiment=EXP
)
DES1.designmatrix(); DES1.FCalc(weights=[0.25,0.25,0.25,0.25])
~~~

Now using matplotlib, we can plot the convolved design matrix:

~~~
import matplotlib.pyplot as plt
plt.plot(DES1.Xconv)

\includegraphics[scale=0.35]{figures/Figure4.pdf}
~~~

We can now define a new design and compare both designs:

~~~
DES2 = neurodesign.design(
  order = [0,0,0,0,0,1,1,1,1,1,0,0,0,0,0,1,1,1,1,1],
  ITI = [2]*20,
 experiment=EXP
)
DES2.designmatrix(); DES2.FCalc(weights=[0.25,0.25,0.25,0.25])
print(“Ff of Design 1: “+str(DES1.Ff))
print(“Ff of Design 2: “+str(DES2.Ff))
print(“Fd of Design 1: “+str(DES1.Fd))
print(“Fd of Design 2: “+str(DES2.Fd))

Ff of Design 1: 0.8571428571428572
Ff of Design 2: 0.4285714285714286
Fd of Design 1: 0.0879554751884
Fd of Design 2: 0.266229261071
~~~

As the second design ignores the presence of the third condition, the frequency optimality (*F*_*f*_) is much worse. However, the blocked character of the design largely improves the detection power. The principles of the genetic algorithm, such as crossover, can be applied to the designs:

~~~
DES3,DES4 = DES1.crossover(DES2,seed=2000)
DES3.order

[0, 1, 2, 0, 1, 2, 0, 1, 1, 1, 0, 0, 0, 0, 0, 1, 1, 1, 1, 1]

DES4.order

[0, 0, 0, 0, 0, 1, 1, 1, 2, 0, 1, 2, 0, 1, 2, 0, 1, 2, 0, 1]
~~~

### 4.4 Generating a random design

The package contains functions to generate random designs. We can generate a random order of stimuli using the function *neurodesign.generate.order*. Below, we gerate a random order of stimuli, which is done by sampling from a multinomial distribution. Below, the resulting probabilities for each trialtype are shown.

~~~
order = neurodesign.generate.order(
   nstim = 4,
   ntrials = 100,
   probabilities = [0.25,0.25,0.25,0.25],
   ordertype = ‘random’,
   seed=1234
)
print(order[:10])
from collections import Counter
Counter(order)

[3, 0, 0, 2, 0, 0, 2, 2, 2, 0]
Counter({0: 36, 1: 22, 2: 22, 3: 20})
~~~

Similarly, we can generate ITI’s from 3 different distributions: fixed (all ITI’s equal), uniform or from a truncated exponential distribution. Below we show the use of the *neurodesign.generate.iti* function and evaluate its output.

~~~
iti,lam = neurodesign.generate.iti(
  ntrials = 40,
  model = ‘exponential’,
  min = 2,
  mean = 3,
  max = 8,
  resolution = 0.1,
  seed=2134
)
print(iti[:10])
print(“mean ITI: %s \n\
  min ITI: %s \n\
  max ITI: %s”%(
    round(sum(iti)/len(iti),2),
    round(min(iti),2),
    round(max(iti),2)))

[0. 2. 2.1 2. 2. 2. 5.4 2. 2.4 5.1]
mean ITI: 2.93
min ITI: 0.0
max ITI: 6.9
~~~

### 4.5 Optimising the design

To optimise the design, we use the *neurodesign.optimisation* class. All parameters are described in Table 2.

~~~
POP = neurodesign.optimisation(
   experiment=EXP,
   weights=[0,0.5,0.25,0.25],
   preruncycles = 10000,
   cycles = 10000,
   folder = “./”,
   seed=100,
   optimisation=‘GA’
   )
POP.optimise()
~~~

**Table 2:** The arguments for object of class *neurodesign.optimisation*

| argument | description |
| --- | --- |
| experiment | The experimental setup of the fMRI experiment (of class `neurodesign.experiment`) |
| G (default = 20) | The size of each generation |
| R (default = [0.4,0.4,0.2]) | The rate with which the orders are generated from (a) blocked designs, (b) random designs and (c) m-sequences |
| q (default = 0.01) | The percentage of mutations in each generation |
| weights | The weights attached to $[F_e, F_d, F_f, F_c]$ |
| I (default = 4) | The number of immigrants in each generation |
| preruncycles | The number of pre-run cycles to find the maximum value of $F_e$ and $F_d$ |
| cycles | The number of cycles in the optimisation |
| seed | The random seed for the optimisation |
| Aoptimality (default = True) | Optimises A-optimality if true, else D-optimality |
| convergence (default = 1000) | After how many stable iterations is there convergence |
| folder | The local folder to save the output |
| outdes (default = 3) | The number of designs to be saved |
| optimisation (default = 'GA') | The optimiser of choice ('GA' or 'simulation') |

## 5 Neurodesign: the GUI

To make the methods more publicly available, we have created a graphical user interface running in a web-application. The back-end of the application is written in python and uses the python module neurodesign described above, the front-end is generated using django, and the application is deployed through a multi-container docker environment on Amazon Web Services.

There are 5 crucial windows of the GUI: main input, contrasts and probabilities, review, console, and settings. The main input window has fields for most parameters from Table 1. Only the parameters P and C are asked in the second window (‘Contrasts and probabilities’). The review window shows all parameters and also prints out the default settings for the genetic algorithm. These parameters, presented in Table 2, can be adjusted in the settings window. The console allows for the optimizations to be started, stopped and followed. When a design optimization is started, the user receives an email with a link to the console where the optimization can be followed (Figure 4). Once the optimization is finished, a zip file can be downloaded containing a chosen number of designs. Each design contains the onsets for each stimulus, a report with design diagnostics (such as collinearity among regressors, see Figure 5), and a script. The script can be used for future reference, or for regenerating the designs locally.

**Figure 4:**
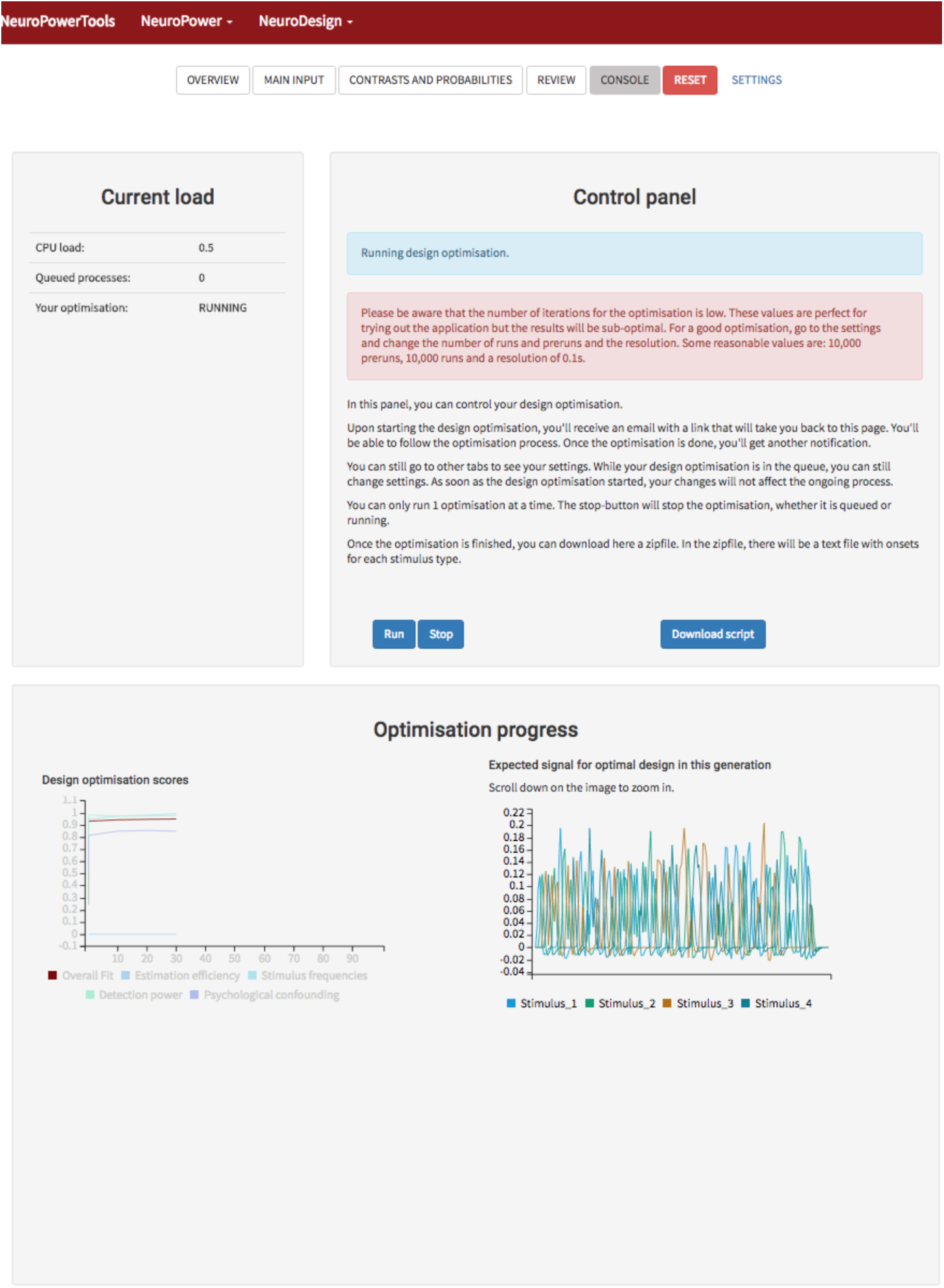
Screenshot of the console where the optimisation can be followed. Every 10 generations, the design is updated with the latest score and the best design.

**Figure 5:**
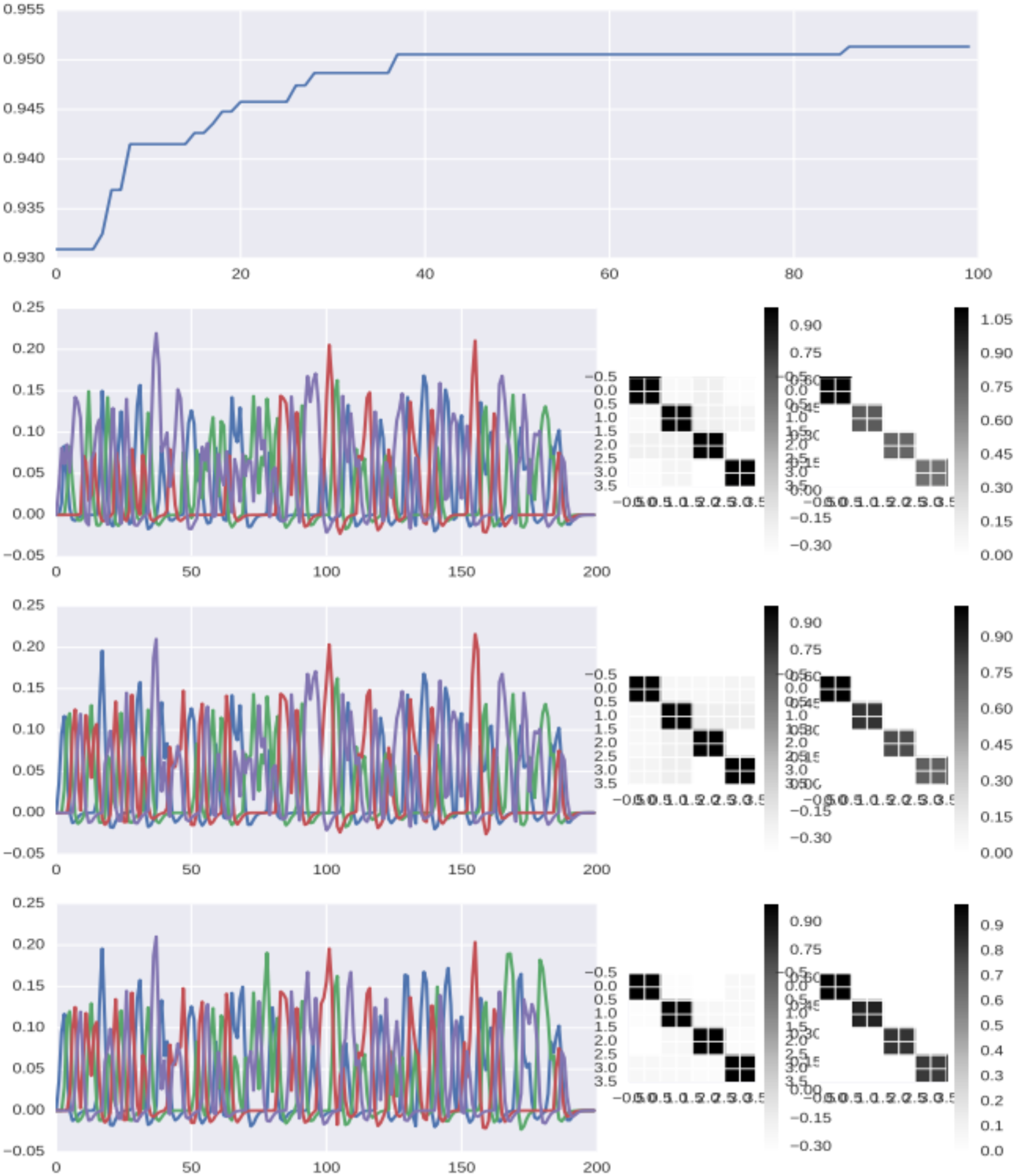
Screenshot of the report describing the optimisation and the best 3 designs from the optimisation.

## 6 Comparison with other software

There are a few alternative design optimisation programs available:

### 6.0.1 mseq

This script, available at http://gru.stanford.edu/svn/matlab/buracas.m was distributed with Buracas and Boynton (2002) and can be used to generate m-sequences using *matlab*.

### 6.0.2 fmri GLM efficiency

This is a tool provided by Henson (2006) to compute the efficiency of experimental designs for fMRI in *matlab* and is available online at https://github.com/MRC-CBU/riksneurotools.

### 6.0.3 mttfmri toolbox

The Multiple Trial Type fMRI MATLAB Toolbox (mttfmri) can generate m-sequences, random designs and blocked designs, as well as compute their efficiency. The toolbox is distributed alongside Liu (2004a) and Liu (2004b). Using the toolbox, a theoretical trade-off between efficiency and power can be calculated. The toolbox assumes a fixed inter-trial interval. The toolbox can be obtained from http://fmriserver.ucsd.edu/tliu/mttfmritoolbox.html.

### 6.0.4 Optseq

*Optseq2* is part of the *Freesurfer* suite (Dale, 1999) and is a *C* toolbox for automatically scheduling events for rapid-presentation event-related experiments. It is available in command line, and it is a simulation-based optimisation. With the toolbox, the order of the stimuli is optimised to be first-order counter-balanced, and random amounts of NULL stimulus are inserted in the design to jitter ITI’s. The total duration and the minimum and maximum NULL time can be specified. Three cost functions can be optimised:

- `eff`: the efficiency: 1*/*trace(*C*(*X*^T^*X*)^−1^*C*^T^)
- `vrfavg`: the average variance reduction factor (VRF): Avg(1*/C*(*X*^T^*X*)^−1^*C*^T^)
- `vrfstd`: a weighted combination of the average and the standard deviation of the VRF’s.

### Selected designs

The following figure shows in the upper panel the optimisation score over the different generations. Below are the expected signals of the three best designs from different families. Next to each design is the covariance matrix between the regressors, and the diagonalmatrix with the eigenvalues of the design matrix.

*Optseq2* by default optimises the design to use the FIR model, but can be customised to optimise the design to use the convolved model.

### 6.0.5 Genetic Algorithm

Wager and Nichols (2003) proposed the use of the genetic algorithm for design optimisation, and published a *matlab* toolbox alongside, available online at http://psych.colorado.edu/tor/Software/genetic_algorithms.html. The application implements a genetic algorithm to optimise experimental designs similar to this implementation. The differences are discussed below.

### 6.0.6 ER-fMRI

*ER-fMRI* is software described in Kao (2009) and implements the genetic algorithm described in Kao et al. (2009). The implementation is similar as the implementation offered by Wager and Nichols (2003) with two key differences: (1) To compute a weighted average *F* of the different optimisation scores, these scores (*Fe, Fd, Fc, Ff*) need to be on a unit scale. While Wager and Nichols (2003) rescale the scores for efficiency (*Fe*) and detection power (*Fd*) in each iteration of the genetic algorithm, Kao et al. (2009) propose to perform a pre-run in which only the *Fe* or *Fd* optimised, in order to povide an optimal score. In other words, Wager and Nichols (2003) rescale the scores within populations, while Kao et al. (2009) rescale the scores over populations. (2) the algorithm starts not only with random designs, but also includes afforementioned the *m*- sequences and blocked designs.

### 6.0.7 Main differences with neurodesign

*Neurodesign* includes all options of all the implementations above: the library can (1) produce *m*-sequences, (2) calculate efficiency scores, (3) generate random and blocked designs, (4) optimise the multi-objective criterion presented in Wager and Nichols (2003) and Kao et al. (2009), as well as (5) optimise designs using the genetic algorithm and a simulation-based optimiser. Whereas the approach of neurodesign is very closely related to *ER-fMRI*, we implement more elaborate control of the ITI’s: in *ER-fMRI* and *Genetic Algorithm*, the ITI’s are modeled simply by introducing null events at random places during the experiment instead of experimental stimulation. This offers control of the minimum and maximum ITI, but it does not control the distribution of ITI’s, as is implemented here. Furthermore, contrary to the other packages, we do not make a distinction between event-related responses and epoch responses. Event-related responses are often seen as a stimulus that results in a direct and short response, while an epoch response represents brain activation over a longer time (for example 2 seconds). Assuming an event-related response in fact equals assuming a response with duration equal to the resolution of the design. Since *neurodesign* supports any stimulus duration, it implicitly covers event-related responses and allows epoch responses. *Neurodesign* also comes with a GUI, which increases the ease of use and accessibility of the application. At the same time, reproducibility of results is ensured by allowing scripting (or downloading scripts) that can be run in a containerised computing environment.

## 7 Design optimisation and statistical power

### 7.0.8 Optimisation

To demonstrate the effect of optimising the experimental design, we performed a simulation study. We compare 3 possible designs: a design optimised using the genetic algorithm (GA), a design optimised using simulations (SIM), and a random design (not optimised - RND). We generated 100 random designs, and report the interval between the 5th and 95th percentile as a non-parametric prediction interval. All designs are generated with *neurodesign*.

We are planning an fMRI study with 3 different stimuli with equal probabilities. The TR is 2 seconds, and there are 450 trials of 1 second each. The ITI’s are sampled from a truncated exponential distribution with (min,mean,max)- values of (0.3, 1, 4) seconds. As such, the total duration of the experiment is 15 minutes, and there are 450 observations. We optimise for 4 different contrasts: [1, 0, 0], [0, 1, 0], [0, 0, 1], [1, 0, −1]. We ran 1000 cycles (pre-run and optimisation). We choose the following values for the multi-objective criterion: *w*_*c*_ = 0.25, *w*_*f*_ = 0.25, *w*_*d*_ = 0.5, since we are mainly interested in statistical power, while keeping the predictability and frequency of the design under control. The evolution of the optimisation scores can bee seen in Figure 6. 90% of the optimisation scores (*F* ^RND^) of the random design are in [0.61 − 0.70], while the resulting scores for the optimised designs are *F* ^GA^ = 0.87 and *F* ^SIM^ = 0.80. The three resulting designs can be seen in Figure 7.

**Figure 6:**
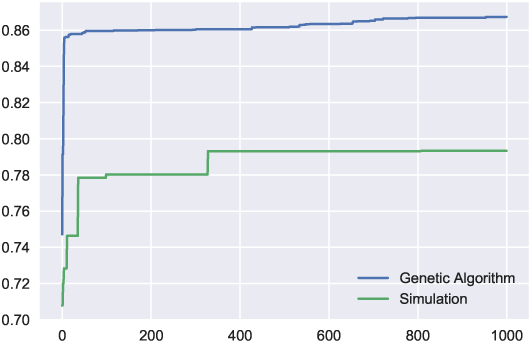
Multi-objective criterion evolution over 1000 iterations for an experimental design with 3 stimuli of 15 minutes.

**Figure 7:**
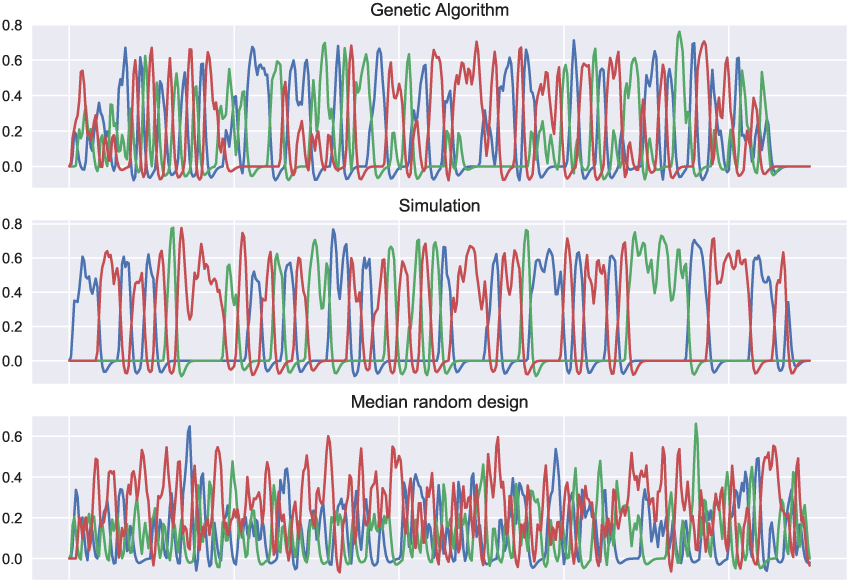
The resulting three designs after optimising using the genetic algorithm, simulation based optimisation, or without optimising (median *F*).

**Figure 8:**
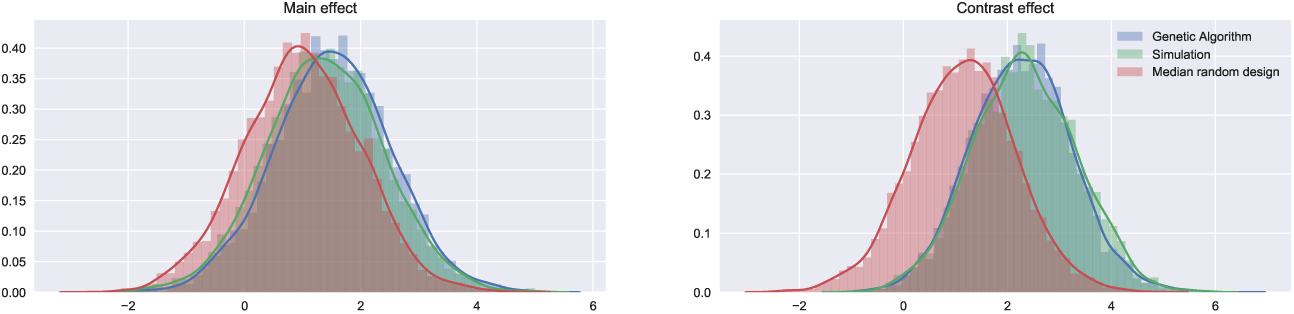
The distributions of *T*-statistics when simulating BOLD signal using the three designs obtained using (1) Genetic Algorithm optimisation, (2) Simulation based optimisation, (3) Random draw (median *F*).

### 7.0.9 Simulation

Based on the three obtained designs, we simulate fMRI data. We simulate data according to equation 1, with *X* the design matrix obtained from the prevous step, *β* = (0.5, 0, −0.5)┬ and *σ* = 1. For simplicity, we ignore the temporal autocorrelation and analyse the data using a simple linear regression model. Specifically, we look at the contrasts (1, 0, 0) and (1, 0, −1), respectively corresponding to absolute effect sizes of 0.5 and 1.0. The distribution of the resulting *T* -statistics based on 1, 000 simulations are shown in 8. We assume a single test, thus performing statistical inference with *α* = 0.95. From the simulations, the observed statistical power is calculated as the percentage a *T* -value exceeds the threshold, #*T > t*_*α*_*/*10^4^. Table 3 shows the resulting observed statistical power. We effectively show how using *neurodesign* significantly increases the statistical power, compared to a random design. Note that even though certain randomly drawn designs result in higher power, these necessarily score lower in the other metrics, such as predictability (*F*_*c*_) and stimulus frequencies (*F*_*f*_), since none of the random designs result in a higher score for the multi-objective criterion.

**Table 3:**
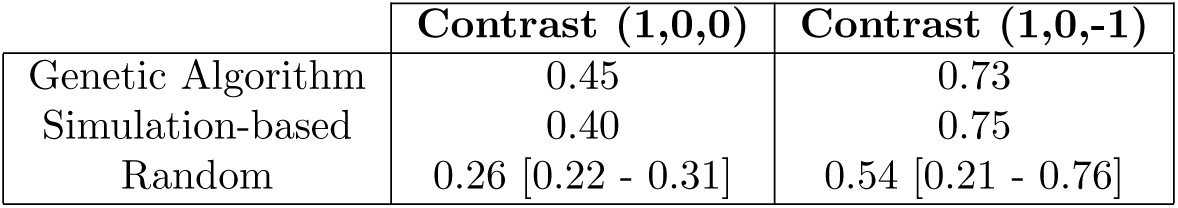
Observed statistical power from 10000 fMRI simulations with 3 different designs: (1) optimised using the genetic algorithm, (2) optimised using simulations and (3) not optimised (median [P5 - P95]).

## 8 Discussion

### 8.1 Default settings

Both the initialisation of the experiment, represented by the class experiment in the python module and the main input window in the GUI, and the genetic algorithm, represented by the class population in the python module and the settings window in the GUI, have some default settings. The default settings of the python module, shown in Tables 1 and 2, are pre-set to ensure a good initial optimisation, while the GUI’s default settings are pre-set for short optimisation duration. While the default settings for the GUI can lead to a sub-optimal design, the user is warned (with a big red textblock at the top of the page) that the settings should be changed if a good optimisation is required. We have chosen these sub-optimal defaults for the GUI to provide a fast run through for first time users, as well as to avoid memory and CPU overload on the server end. For the experiment, we assume a priori that there are no rest blocks and that the trial only consists of stimulation (no fixation cross etc.). There is by default no limit on the maximum number of times a stimulus can be repeated, and the stimulus frequency is not controlled with a hard limit. The default resolution in the python module is 0.1 seconds, while in the GUI it is 0.25 seconds. For the genetic algorithm, the default settings are as follows. The optimisation calculates the A-optimality. In each generation, the percentage of mutations is 1%, the number of immigrants is 4 designs and the size of each generation is 20 designs, as is suggested by Kao (2009). When generating new designs, there are 40% blocked designs, 40% random designs and 20% m-sequences. Convergence is reached when the score is stable for 1000 generations. There are no default settings on the number of cycles in the python module, while the GUI runs by default 10 cycles to find the maximum *F*_*d*_ and *F*_*e*_ and 100 cycles for the optimisation (again with a clear message that this can be increased for more optimal results).

### 8.2 Reproducibility

In line with the recent effort to make neuroimaging research fully reproducible, this application makes it possible to track the exact source of each design. Low level reproducibility is provided by making a script available for download with which the optimisation can be regenerated. Running this script in python, given that the required libraries are installed, will repeat the analysis. However, this script will repeat the analysis but does not guarantee the same results as the specific configuration of the computer on which the analysis is run can influence the results.

Higher level reproducibility, that guarantees replicability not only of the analysis but also of the results, is possible with the use of Docker containers, which is a small piece of software that emulates a given computational configuration (operating system, libraries, python packages,…). Based on the model presented by BIDS-apps (Gorgolewski et al., 2017), our analyses run in Docker containers that are open-source and available for download at https://hub.docker.com/r/neuropower/neuropower Running the following command in a terminal will replicate the analysis that has been performed through the GUI.

~~~
docker run -v /location_where_the_script_is/:/local \\
-it neuropower/neuropower python /local/name_of_the_script.py
~~~

This use of Docker containers is not only well suited for reproducibility of the GUI, but also allows the replication of results from a python script (given that a random seed is set).

## 9 Conclusion

We present a toolbox for optimizing fMRI designs. The toolbox is an extension of currently available toolboxes, allowing for more complex design and better control and optimization of timing of stimuli. The toolbox is available through different modalities: a user-friendly GUI accessible at www.neuropowertools.org and a python package. The code is available on www.github.com/users/neuropower.

## 10 Acknowledgements

This work was supported by the Laura and John Arnold Foundation. J.D. has received funding from the European Unions Horizon 2020 research and innovation programme under the Marie Sklodowska-Curie grant agreement No 706561.

